# Concurrent fMRI demonstrates propagation of TMS effects across task-related networks

**DOI:** 10.1101/2022.01.06.475293

**Authors:** Lifu Deng, Olga Lucia Gamboa, Moritz Dannhauer, Anshu Jonnalagadda, Rena Hamdan, Courtney Crowell, Tory Worth, Angel V. Peterchev, Marc A. Sommer, Roberto Cabeza, Lawrence G. Appelbaum, Simon W. Davis

## Abstract

Transcranial magnetic stimulation (TMS) has become an important technique in both scientific and clinical practices, and yet our understanding of how the brain responds to TMS is still limited. Concurrent neuroimaging during TMS may bridge this gap, and emerging evidence suggests widespread that modulatory effects of TMS may be best captured through changes in functional connectivity between distributed networks, rather than local changes in cortical activity. However, the relationship between TMS stimulation parameters and evoked changes in functional connectivity is unknown. In this study, 24 healthy volunteers received concurrent TMS-fMRI while performing a dot-motion direction discrimination task. An MR-compatible coil was used to apply trains of three pulses at 10 Hz rTMS over the primary visual cortex (V1) at the onset of the dot stimuli with four levels of stimulation intensity (20%, 40%, 80%, and 120% of resting motor threshold, RMT). Behavioral results demonstrated impairment of motion discrimination at 80% RMT. FMRI results yielded three findings. First, functional connectivity between visual and non-visual areas increased as a function of rTMS intensity. Second, connectivity *within* the visual network was positively associated with motion accuracy, while the connectivity *between* visual and non-visual regions was negatively associated with motion accuracy. Lastly, we found that reductions in the similarity between functional and structural connectivity associated with increasing TMS intensity were constrained to the visual network. These findings demonstrate spatially dependent nonlinear effects of TMS intensity on brain functional connectivity that proceed beyond the site of stimulation and influence associated behavior.

## 1. INTRODUCTION

Transcranial magnetic stimulation (TMS) has been a valuable tool in both cognitive and clinical neuroscience because it allows for the examination of specific neurocognitive mechanisms and the development of effective therapies based on these mechanisms. Ideally, the understanding of mechanisms should precede clinical application, but even the simple dose-response relationships of TMS are largely unknown outside of the motor cortex. Multiple lines of evidence in animal models and human studies suggest that TMS induces complex changes in activity across broad networks of cortical regions, demonstrating clear evidence of polysynaptic effects (Wang and Voss, 2015; Cocchi et al., 2016; Alekseichuk et al., 2019; Beynel et al., 2020b). In line with these empirical findings, a recent systematic review of TMS on resting state networks (Beynel et al., 2020a) found that while network-level effects were common in the literature, these effects did not follow the typical frequency-dependent heuristic that helps to predict the local effect of TMS modulation (Luber and Deng, 2016). As such, it can be inferred that network-level modulatory effects depend on not only the parameters of stimulation but also the functional interactions between the stimulated and connected regions.

TMS during concurrent neural recordings has become an exciting technological development that may expand the repertoire of addressable questions by leveraging the rich spatial and temporal information afforded by different imaging methods. Previous studies utilizing concurrent TMS and fMRI are often focused on resting-state brain activity without the presence of cognitive tasks. In such studies, stimulation of primary sensory or motor cortices frequently is reported to exert feed-forward influences on functionally relevant regions, such as the frontal eye fields (Cocchi et al., 2016; Castrillon et al., 2020). Similarly, concurrent TMS during electroencephalographic (EEG) recordings has shown that single-pulse TMS to the dorsal attention network results in widespread activation of that same network (Ozdemir et al., 2020). There is also evidence that these intra-network effects are facilitated by structural connectivity (Momi et al., 2021a), suggesting that stimulation propagates between brain regions that are structurally connected to the stimulation site.

A lingering question is the relevance between these TMS-driven dynamics and cognition, as the effects of TMS has been shown to depend on the cognitive state (Silvanto et al., 2018). For example, when rTMS is applied to motion selective cortex (MT+) during a motion discrimination task, performance is hindered when attention is focused on motion but enhanced when attention is focused on other visual attributes (Brascamp et al., 2010; Zinchenko et al., 2021). Relatedly, connectivity can depend on brain state during motion perception, based on evidence that frontal and occipital connectivity is only present when participants are aware of apparent motion (Sanders et al., 2014). In addition, the timing of stimulation relative to the onset of visual motion strongly determines the influence on performance and EEG responses (Gamboa Arana et al., 2020), implying the different cognitive components affected. Because attention-driven activity during motion-perception tasks is influenced by gating between primary visual cortex and MT+ (Stephan et al., 2008), a network approach may better explain the effects of TMS than focusing on a single region.

Building on past studies on concurrent TMS-neuroimaging, we sought characterize the effects of TMS on network-level functional connectivity (FC) due to different intensities during visual motion perception. Graph theoretical analyses were used to depict the organization of functional sub-systems and how TMS influenced communication between these systems. Three specific goals were pursued. First, this study investigated whether TMS affected global or local FC, and whether the effects were constrained to the network related to the task. Secondly, this study examined the effects of FC on motion discrimination task performance to better understand how changes in FC mediate behavior. Lastly, structural connectivity (SC) was used to explore how TMS alters the functional network relative to the underlying architecture. This analysis combines structural, functional, and behavioral information to further the understanding of the impacts of TMS on global brain network and behavior.

## 2. METHODS

### 2.1 Subjects

A total of 27 healthy volunteers (16 females, mean age = 23.7 years, SD= 2.78) participated in this study. At the beginning of the first visit, all participants were informed about the experimental procedures and were screened for contraindications to MRI and TMS. Subsequently, written informed consent was obtained from each participant. All participants were right-handed according to the Edinburgh Handedness Inventory (Oldfield 1971), had normal or corrected-to-normal vision, and did not have a history of neurological or psychiatric diseases. Participants received $20 per hour for taking part in this study. Procedures in this study conformed to the Declaration of Helsinki and were approved by the Institutional Review Board of Duke University. Data from three participants were excluded from the study as they declared falling asleep during the MRI session. As such, the final dataset analyzed for this study consisted of 24 healthy volunteers (13 females, mean age = 23.6 years, SD= 2.55).

### 2.2 Experimental design

Participants in this study took part in two experimental sessions. During the first session, participants were provided with screening questionnaires and were administered drug and pregnancy tests to check for eligibility. Resting motor threshold (RMT) was then acquired and subjects were briefly familiarized with the motion discrimination task by performing 1-2 minutes of practice trials. During the second session, participants first practiced the motion discrimination task in a staircase fashion inside a mock MRI scanner that mimics the environment of real MRI scanner (projection system, response box, recorded scanner sound, etc.). Before starting the fMRI-TMS data collection, participants were positioned in the bore of the MRI scanner and performed one additional staircase of the motion discrimination task to determine the threshold coherence level used in the subsequent fMRI-TMS task, as described in *Section 2*.*3*.*1*. After fMRI-TMS data acquisition, additional structural and diffusion weighted images (DWI) were acquired.

### 2.3 Procedures

#### 2.3.1 Motion perception task

Stimuli were generated using MATLAB (Mathworks, Natick, MA, USA) and the Psychtoolbox extension (http://www.psychtoolbox.org). Each trial started with a white fixation-cross shown in the middle of the screen on a black background (see **Fig. 1A**). After 530 ms, white dots (5 pixels each, dot density: 1.41 dots/deg^2^, dot- speed: 7.4 °/s) were presented for 150 ms, moving at a pre-determined coherence level (see next paragraph) either upwards or to the left within a circular window on the upper right quadrant of the screen. The center of the dot field was located at 8° from the fixation cross and extended 12° in diameter. Stimuli were projected onto a mirror placed above the scanner head coil using a rear projection system. The projector refresh rate was 60 Hz and viewing distance between the mirror and the projector screen was 70 cm. When needed, vision was corrected using MRI-compatible lenses matching a participant’s prescription. Responses were recorded via a fiber-optic response box (Resonance Technology, Inc). Participants had up to 2 s to indicate if dot motion was to the left using the right index finger or upwards using the right middle finger. After each response, a cross colored either green (correct response) or red (incorrect response) was shown during 1 s to provide feedback. Participants were instructed to perform the task as accurately and as quickly as possible, with an emphasis on accuracy.

**Figure 1.**
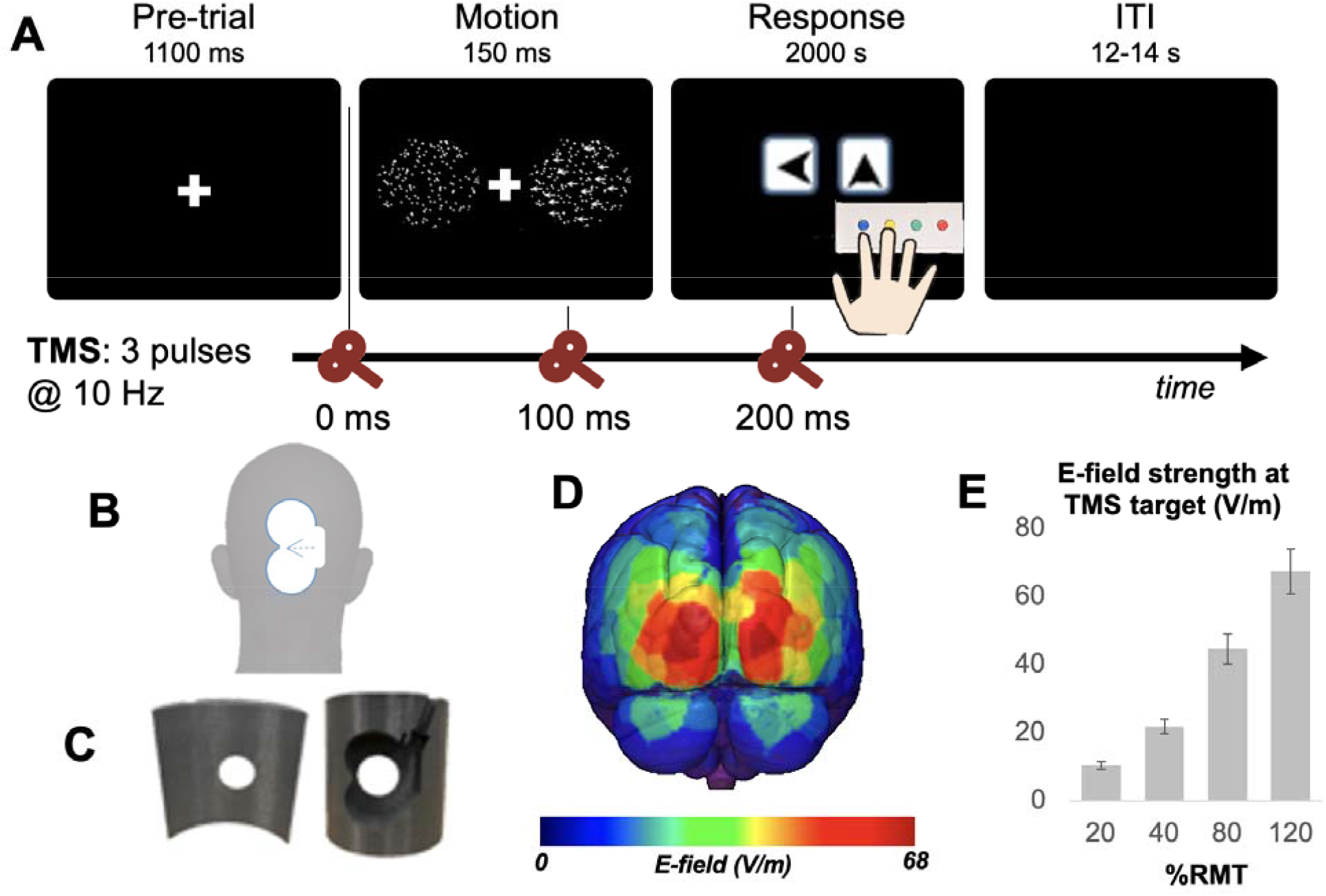
Experimental design. **(A)** Illustration showing still frames of the visual stimulus along with three TMS pulses (orange markers) presented at 10 Hz at the start of stimulus presentation. **(B)** Coil orientation and stimulation site over Oz as defined in the 10-20 EEG System. **(C)** 3D-printed headrest used in the MRI scanner to position the TMS coil over the stimulation site on Oz. Four different intensities were used: Oz stimulation at 20, 40, 80 and 120% RMT as the main tests and vertex stimulation at 120% RMT as control. **(D)** Simulation of the TMS electric field (E-field) strength induced within each cortical node, averaged over all participants. **(E)** Simulated E-field strength (mean ± standard deviation) at the target cortical node for each Intensity level across participants.

Before starting the TMS-fMRI data collection during the second session, subjects performed a final practice version of the motion perception task within the bore of the magnet comprising 120 trials in which dot coherence varied from 0% coherence to 100% coherence according to a staircase procedure. Data from this practice session was assessed online to ascertain an individualized coherence value to use in the main task. As in our previous work with this paradigm (Gamboa Arana et al., 2020; Gamboa et al., 2020), the calculation was done by means of a generalized linear model (MATLAB’s glmfit function with ‘binomial’ option) that fit a sigmoid function to the signed (+1/−1 due to dot motion direction) trial coherences and individual correct/incorrect responses to find a particular coherence where 70% of the training trials were correct.

#### 2.3.2 Concurrent TMS–MRI

After the practice task in the mock MRI scanner, subjects were transferred to the MRI scanner room. Participants wore a tightly fitting swimsuit cap where the stimulation points were marked. The main stimulation target was centered over Oz as defined in the 10-20 EEG system, which is located 10% of the distance between the inion and nasion (approximately 3 cm above the inion in most adults, see **Fig. 1B**) (Ishikawa et al., 2011). The coil was placed inside a custom headrest that was 3D printed from semi-flexible material (nGen_FLEX) to be comfortable for the subject and maintain the TMS coil position on the head during pulses (**Fig. 1C**). The headrest was specially designed for each stimulation target so that the center of the coil was tangential to the stimulation point and the induced electric field was perpendicular to the midline, while the cable-side of the coil was on the right hemisphere. Positioning was adjusted both before the scan using markers on the fitted swim cap and was verified after the experiment with the use of four MR-sensitive fiducial markers placed over the borders of the major and minor semi-axes of the coil. Cushions were used to reduce head motion inside the scanner and participants were informed about the importance of minimizing head motion during task performance. Participants wore earplugs to attenuate scanner noise and the TMS clicking sound.

A burst of three biphasic TMS pulses at 10 Hz delivered at motion onset was applied over the stimulation site using a MagPro R30 magnetic stimulator connected to an MRI-compatible TMS figure-of-eight coil (MRi-B91 Air-cooled, MagVenture, Farum, Denmark). Stimulation over the visual area was delivered at 20%, 40%, 80% and 120% of the participant’s RMT, presented in pseudo-randomized order in each of the runs. The task was presented in the form of a slow event-related design, such that the duration of each trial was 15 s. The TMS device, the presentation computer, and the MR acquisition were synchronized via an Arduino board (**Fig. 2A**), such that the TMS pulses were delivered reliably in between slice acquisitions. Furthermore, timing of the MR acquisition rate (30 slices every 2 seconds, or 15 Hz) and TMS frequency (10 Hz) allowed us to place the TMS pulses consistently inferior (slice 4/30, roughly at the level of the lower cerebellum) and superior (slice 28/30) locations, ensuring that artifacts associated with the TMS pulse were spaced at least 4–6 slices away from the principle regions of interest (i.e., visual cortex, **Fig. 2B**); Arduino code used to time pulses can be found online in our public repository (https://github.com/ElectricDinoLab). The task was divided into five runs, with each run containing 32 trials (8 trials for each intensity), giving a total 40 trials per Intensity over the course of an fMRI session. As each trial comprised three 10 Hz pulses, a total of 120 pulses were administered at each intensity.

**Figure 2.**
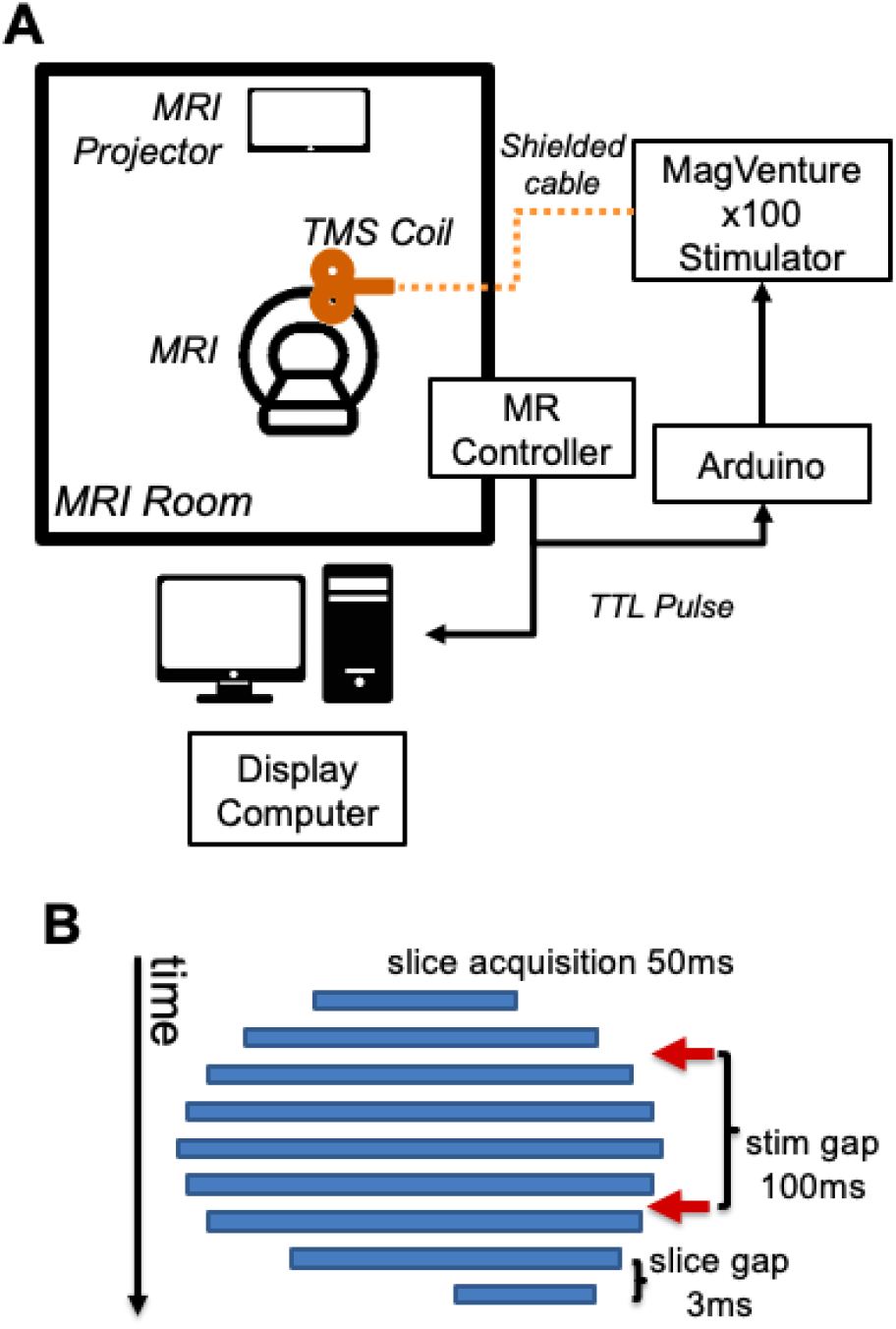
Concurrent TMS-fMRI. **(A)** A generalized wiring diagram. **(B)** Schematic of the timing of TMS pulses to avoid concurrent acquisition of data and TMS pulses.

#### 2.3.3 Resting motor threshold (RMT) measurement

To determine the TMS intensities to use during the experiment, the resting motor threshold (RMT) of the right first dorsal interosseous muscle (FDI) representation in the left primary motor cortex was obtained in the first visit. During this procedure, biphasic single pulses were sent with a MagPro R30 stimulator connected to a figure-of-eight coil (MCF-B65 cool coil, MagVenture, Farum, Denmark) while being guided by a stereotaxic neuronavigation system (Brainsight, Rogue Research, Canada). Surface electromyogram (EMG) was recorded from the right FDI muscle by using disposable electrodes (Covidien / Kendall, 133 Foam ECG Electrodes, Mansfield, USA) in a belly tendon montage. RMT was defined as the minimum intensity at which a motor evoked potential (MEP) of 50 μV average peak-to-peak amplitude was elicited when the muscle was at rest (Conforto et al., 2004). The Motor Threshold Assessment Tool software (MTAT 2.0) (http://www.clinicalresearcher.org) was used to determine the RMT once a reliable stimulation site for the FDI motor ‘hot spot’ was found. The percentage of the maximum stimulator output and the realized peak coil current rate of change (dI/dt) values were registered to be used in the following session.

#### 2.3.4 fMRI data acquisition & analysis

Functional MR images were obtained with a 3T GE MRI system scanner with a 4-channel transmit/receive head coil at the Brain Imaging and Analysis Center (BIAC) at Duke University Medical Center. During the visual task an echoplanar imaging (EPI) sequence was used (320 volumes, 30 axial slices, repetition time/echo time =1500/27 ms; flip angle = 70; FOV = 256 mm^2^; matrix size = 64×64; voxel size = 4×4×4 mm^3^; slice thickness = 4mm). The first 9 s (6 volumes) of the fMRI record were discarded to allow for T1 equilibrium magnetization. Anatomical images for co-registration were collected using a T1-weighted MP-RAGE sequence (repetition time/echo time/inversion time = 6.972/3.01/400 ms, flip angle =11º).

Functional image processing proceeded first by removing slices affected by TMS pulses. This was necessary because, even though TMS pulses were timed to be delivered in between MR slice acquisitions, small errors in timing may have resulted in overlap between the TMS pulse and MR acquisition. We identified slices with a signal magnitude of > 2 SD from the run mean and visually inspected them for the presence of the TMS artifact. These slices were replaced by temporal interpolation of the signal values of the same slice from the preceding and succeeding volumes. Functional images were then preprocessed using image processing tools, including FLIRT (FMRIB’s. Linear Image Registration Tool) and FEAT (FMRIB Expert Analysis Tool) from FMRIB’s Software Library (FSL, https://fsl.fmrib.ox.ac.uk/fsl/fslwiki/). Images were corrected for slice acquisition timing, motion, and linear trend; motion correction was performed using FSL’s MCFLIRT, and six motion parameters estimated from the step were then regressed out of each functional voxel using standard linear regression. Images were then temporally smoothed with a high-pass filter using a 190 s cutoff and normalized to the MNI stereotaxic space. To remove any additional artifacts associated with TMS pulses, spatiotemporal independent components analysis (ICA, FSL’s MELODIC) was used to identify and remove noise components occurring in slices collected concurrent with the TMS pulse. White matter and CSF signals were also removed from the data and regressed from the functional data using the same method as the motion parameters. BOLD activations during the initial dot motion presentation were entered in a standard GLM, using HRF-convolved trial regressors and their temporal derivatives.

#### 2.3.5 Electric field modeling

The direct effects of TMS are driven by the strength and spatial distribution of the electric field (E-field) induced in the brain. Thus, simulation of the E-field induced by TMS in individual participants is increasingly recognized as an important step in spatial targeting of specific brain regions and forms a critical link between the externally applied TMS parameters and the neurophysiological response supporting cognitive operations (Peterchev et al., 2012; Gomez et al., 2020). To determine the E-field induced by TMS in the brain of each participant, simulations using T1 images and the finite element method in the SimNIBS software package (version 2.0.1, Thielscher et al., 2015) were conducted (Beynel et al., 2019; Beynel et al., 2020c; Beynel et al., 2021). Twenty-three subjects were included in the E-field analysis and head models were created using the SimNIBS mri2mesh pipeline, featuring five distinct tissue types: skin, skull, cerebrospinal fluid, gray matter, and white matter. The scalp, skull, and cerebrospinal fluid tissue compartments were modeled with isotropic conductivity values of 0.465, 0.01, and 1.654 S/m, respectively. For seventeen subjects, available DWI information was used to estimate anisotropic conductivities for the brain matter using the volume-normalized approach (Gullmar et al., 2010) which kept the geometric mean of the conductivity tensor eigenvalues for the gray and white matter equal to literature-based isotropic values of 0.275 and 0.126 S/m, respectively. DWI data could not be processed for six subjects, for which the latter isotropic conductivity values for gray and white matter were assigned to their corresponding computational tetrahedral elements. For nine subjects, 4 fiducial markers implemented with gel-filled stickers were attached on the active side of the TMS coil to image the location of the coil relative to the head using additional T1 scans. Based on the location of the fiducial markers, the mean (+/- standard deviation) shortest distance between the TMS coil center and the scalp was determined to be 3.8 mm (+/- 3.4 mm) across the nine subjects, and the coil plane deviated from the local scalp tangential plane by only 7.2° (+/- 3.2°), confirming good coil placement. For fifteen subjects it was not possible to consistently image the fiducial markers. Instead, the target Oz scalp site was identified from the MRI data and head model’s scalp surface, the coil center was extruded by 4 mm to account for the scalp–coil spacing, and the coil plane was oriented tangentially to the closest scalp surface location. The individual E-field was simulated for TMS coil current rate of change (dI/dt) of 1 A/μs. Leveraging the direct proportionality between the coil current and the E-field strength, the latter was obtained by scaling the E-field solution for the various TMS intensities based on the corresponding dI/dt value computed from the Intensity setting according to the formula dI/dt = 1.9308 × %MSO – 2.4018. The formula was derived by linear regression of the dI/dt values reported by the TMS device for Intensity settings as percentage of maximum stimulator output (%MSO) for 5% MSO steps averaged over three TMS pulses. **Figures 1D** and **1E** present, respectively, a map of the cortical-node E-field strength at 120% RMT averaged across the subjects, and the E-field strength at the target cortical nodes across the four TMS intensity levels.

#### 2.3.6 Network Analysis

##### Functional connectivity (FC) network construction

A consistent parcellation scheme across all subjects and all modalities (DWI, fMRI) was used. Subjects’ T1-weighted images were segmented using SPM12 (fil.ion.ucl.ac.uk/spm/software/spm12/), yielding a GM and white matter mask in the T1 native space for each subject. The entire GM was then parcellated into 397 ROIs, each representing a network node by using a sub-parcellated version of the Harvard-Oxford Atlas (Fornito et al., 2010). Functional connection matrices representing task-related connection strengths were estimated using a correlational psychophysical interaction (cPPI) analysis. Briefly, the model relies on the calculation of a PPI regressor for each region, based on the product of that region’s physiological time series and a task regressor convolved with the HDR, to generate a term reflecting the psychophysical interaction between the seed region’s activity and the specified experimental manipulation. The psychological regressor was multiplied with two physiological time courses for regions i and j, producing two psychophysiological interaction regressors, denoted as PPI_i_ and PPI_j_. The partial correlation (ρ_XY·Z_, where X= PPI_i_, Y=PPI_j_), was then computed as the partial correlation between two psychophysiological interaction regressors for a given pair of regions *i* and *j*, controlling for the covariates (denoted as *z*, which includes the HDR-convolved task regressor and the BOLD time courses for regions *i* and *j*); the six motion parameters, white matter signal, and CSF signal were also included as confounds.

##### Modularity

Functional sub-network community assignment was identified using modularity analysis (Reichardt and Bornholdt, 2006). A common whole-brain network was obtained by averaging all functional networks across participants and intensity levels. This common network was then thresholded to preserve 15% strongest connections and submitted to the modularity algorithm (gamma = 1). Based on module assignments, functional connections were classified into between-module connections (connecting regions in different modules) and within-module connections (connecting regions in the same module).

##### Structural connectivity (SC) network construction

Information on the structural connections based on diffusion tractography between each pair of regions in the data was assessed with a standard DWI processing pipeline used previously in our group (Davis et al., 2019). DWI data were analyzed using FSL (fsl.fmrib.ox.ac.uk) and MRtrix (mrtrix.org) software packages. Data were denoised, corrected with eddy current correction, and bias-field corrected. Constrained spherical deconvolution was used in calculating the fiber orientation distribution; after tracts were generated, they were filtered using spherical-deconvolution informed filtering of tractograms (SIFT; Smith et al., 2013). To generate the SC network, the value of SC was defined as the number of streamlines connecting each pair of ROIs.

#### 2.3.7 Statistical Analysis

Statistical analyses were performed using MATLAB and RStudio. The effect of TMS intensity on behavior was assessed using repeated-measures ANOVA. The FC network of each participant under each rTMS intensity was first normalized into z-scores to further control for motion-induced confound. The between-and within-module FCs were then derived from the z-scored network. The effect of TMS on FC within and between network was assessed using two-way repeated-measures ANOVA, in which the Type of FC (between/within) and the Intensity of TMS (20/40/80/120% RMT) were defined as the two main effects. The relationship between connectivity and task performance was examined in mix effect regression model using the lme4 package in R, in which the dependent variable was the correct response rate. The random effect was specified as participant-specific intercept. Variables of interests, estimated as fixed effects, were TMS intensity, between-module and within-module FC of the visual module, as well as participant-level covariates such as sex, coherence of the visual stimuli, and resting motor threshold. The neural effects on performance were assessed using F-contrasts and T-contrasts. Details about the design matrix and the contrasts can be found in **Figure S1a** and **Figure S1b**, respectively, in Supplementary Materials.

## 3. RESULTS

### 3.1 Behavioral Results

The accuracy of motion discrimination task (leftwards vs. upwards motion) during concurrent TMS-fMRI was analyzed to assess the behavioral effect of TMS targeting visual cortex. A one-way repeated-measures ANOVA revealed significant differences in performance across TMS intensity levels (SS_subjects_ = 0.69, SS_condition_ = 0.04, F_3,23_ = 2.93, *p* = 0.039). Post hoc, uncorrected comparisons between TMS Intensity conditions revealed that motion accuracy in the 80% RMT condition (mean: 58% ± 9%) was significantly lower than 20% RMT (mean: 63% ± 13%, *p* = 0.011), 40% RMT (mean: 62% ± 10%, *p* = 0.024), and 120% RMT (mean: 61% ± 9%, *p* = 0.047) conditions (**Fig. 3A**). No differences in discrimination accuracy were seen between the highest (120% RMT) and lowest (20% RMT) stimulation intensities (p > 0.05), suggesting that the disruptive effects were attributable to a neural response in visual processing regions rather than distraction-related confounds (sound, somatic sensations) associated with increasing TMS intensity. Reaction times (**Fig. 3B**) were not significantly affected by the different TMS intensities, though correct trials were significantly faster than incorrect trials (paired t_23_ = 5.33, p = 0.0002).

**Figure 3.**
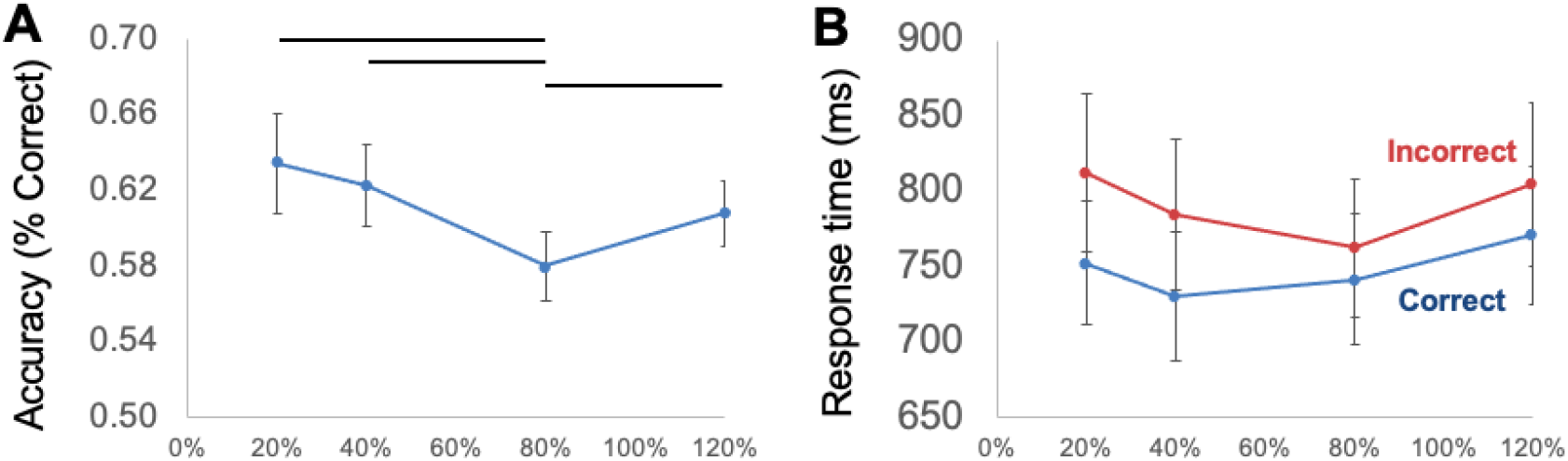
Behavioral results. Accuracy **(A)** and reaction times **(B)** for all participants in the dot-motion perception task during TMS-fMRI. TMS selectively disrupted motion performance at 80% RMT, relative to all other stimulation intensities. In contrast, no effects of Intensity were seen on reaction time.

### 3.2 Delineating task network nodes affected by TMS

The present analysis focused on evaluating network-level, fMRI-based functional connectivity to characterize the effects of stimulation intensity. The network-level effects of TMS were examined on two scales: brain modules, and directly stimulated nodes in visual cortex. Stimulated nodes were defined using E-fields estimated within each individual, localized using fiducial markers placed on the TMS coil; E-fields were then scaled by individual RMT, and then voxelwise E-field values were averaged within a parcellated region of interest to identify the network node with the highest E-field (**Fig. 1D**). Whole brain network data were delineated into meaningful networks to establish reliable boundaries between local and global communities of regions, termed “modules” (also known as “subnetworks”), by using modularity analysis on the average FC matrix (averaging across subject, TMS intensity, and task success conditions). This data-driven modularity analysis parsed 397 ROIs into four highly recognizable modules (**Fig. S2**), including (i) default mode network (DMN), (ii) somatosensory-motor network (SM), (iii) a visual-temporal network complex, and (iv) frontal-parietal control network (FPCN). To obtain a more accurate delineation of the visual system in the Visual-Temporal complex, an additional modularity analysis was performed on it, which in turn produced four subdivisions (**Fig. S3**); among these subdivisions, the two posterior subdivisions were combined into the Visual network (Vis.), and the two anterior, the Temporal network (Tmp.). The final 5-module assignment is shown in **Figure 4A**.

**Figure 4.**
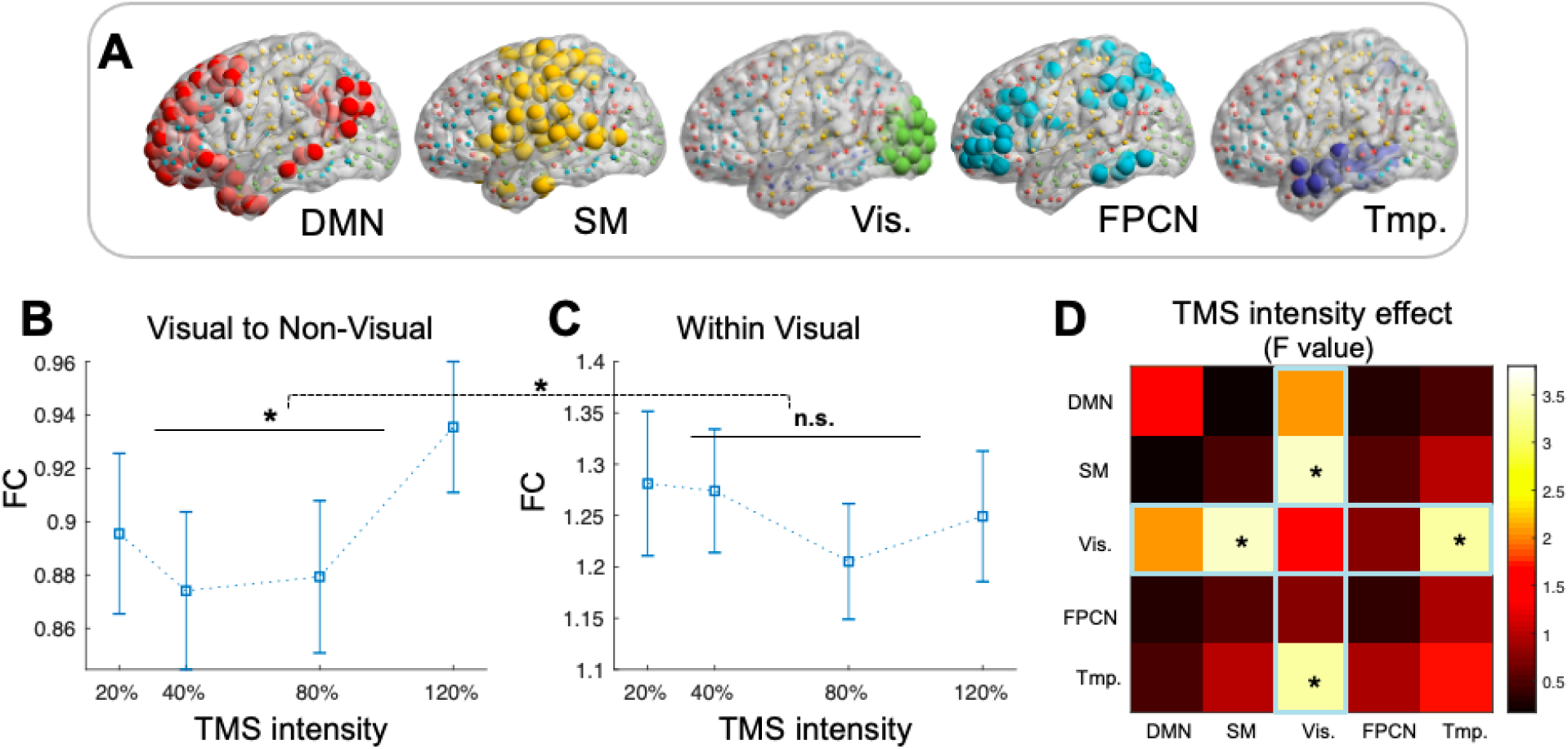
Within and between-module connectivity in response to TMS. (**A)** The 5 modules obtained from modularity analysis (DMN = default mode network, SM = somatosensory-motor network, Vis. = visual network, FPCN = frontal-parietal control network, Tmp. = temporal network). (**B)** and (**C)** The effects of TMS intensity on visual network FC (z-scored); dashed horizontal brace indicates the Intensity Type interaction; horizontal solid lines indicate the main effects of Intensity. (**D**) The effects of TMS intensity (expressed as F-values) on the FC between different modules. Error bars indicate standard error of the mean (SEM). Asterisks (*) indicate p<0.05.

### 3.3 TMS effects on functional connectivity

#### 3.3.1 TMS effects on visual system functional connectivity

Network-level TMS effects were first evaluated on the task-relevant regions. The 5-module community assignment (**Fig. 4A**) was used to define within-module FC (the average FC within the same module) and between-module FC (the average connectivity between the given module and other modules). For each of the five modules, an Intensity (% RMT) by Type (within/between module connectivity) repeated-measures ANOVA was performed. For all five modules, the within-module FC was significantly stronger than the between-module FC (DMN: F_1,21_ = 11.8, p = 0.0023; SM: F_1,21_ = 40.52, p=1.7e-6; Visual: F_1,21_ = 33.04, p = 7.4e-6; Temporal: F_1,21_ =17.45, p = 3.6e-4; FPCN: F_1,21_ = 17.23, p = 3.9e-4), suggesting that the data-driven modularity analysis successfully captured meaningful modules of the whole-brain functional network. For completeness, results for the larger Visual-Temporal complex are shown in **Fig. S2** in the Supplementary Materials, which were similar to the findings for the Visual network.

Crucially, a significant Intensity x Type interaction was found uniquely in the Visual network (F_3,69_ = 3.99, p = 0.011, **Fig. 4B, 4C**) but not in the other modules (all p > .05), suggesting that the TMS may have a type-specific effect on the functional communication of the targeted visual system. The effects of TMS intensity on these two types of FC were further examined in two separate one-way repeated measure ANOVAs, where the intensity effect was significant for between-module connectivity (i.e., ‘Visual to Non-Visual’, F_3,69_ = 2.88, p = 0.039, **Fig. 4B**) but insignificant for within-module connectivity (i.e., ‘Visual to Visual’, F_3,69_ = 1.69, p = 0.178, **Fig. 4C**). An exploratory analysis suggested that such significant modulation of between-module FC seemed to be driven by the connections with SM and Temporal modules; both findings are consistent with the idea that the effects of increasing TMS elicited greater network activity due to increasing motor (SM) or auditory (Temporal) sensation. No significant effect of TMS intensity existed between other modules (**Fig. 4D**). In sum, these results showed that TMS modulated FC of the targeted functional system.

The above results support the idea that intensity effects can be seen at the level of the stimulated network, but a more rudimentary concern might be that these same effects can also be seen at the level of the directly stimulated region. To examine this possibility, two non-overlapping groups of visual ROIs were further investigated: 1) three ROIs in the left occipital pole that were closest to the TMS coil (Group 1, **“directly-stimulated visual cortex”**), which also received visual inputs from the right visual field and, and 2) the ROIs in visual network not directly stimulated (Group 2, **“Indirectly-stimulated visual cortex”**). For each of these two ROI groups, the average FC to other Visual module ROIs (i.e., ‘Within Visual FC’) and the average FC to non-Visual ROIs (i.e., ‘Group 1/2 to Non-visual FC’) were measured. Results largely followed similar, but weaker patterns as found in the above paragraph. For *directly-stimulated* visual cortex (Group 1), the Intensity by Type ANOVA did not produce a significant interaction (F_3,69_ = 1.85, p = 0.15), and two separate one-way ANOVA showed no effect of Intensity in either “Group 1 to Visual” FC (F_3,69_ = 1.18, p = 0.33) or “Group 1 to Non-Visual” FC (F_3,69_ = 0.26, p = 0.85). In comparison, the results for *indirectly-stimulated* visual cortex (Group 2) were largely consistent with the results based on the whole Visual network (**Figs. 4B, 4C**): the Intensity by Type ANOVA revealed a significant Intensity x Type interaction (F_3,69_ = 3.12, p = 0.032), and two separate one-way ANOVA showed no effect of Intensity in “Group 2 to Visual” FC (F_3,69_ = 1.65, p = 0.17) but significant “Group 2 to Non-Visual” FC (F_3,69_ = 3.38, p = 0.023). In summary, these ancillary findings demonstrated the distributed nature of TMS effects across the visual system, with changes in FC largely observed in brain regions that were not directly stimulated by the TMS E-field.

#### 3.3.2 Behavioral correlates of visual system FC

A second goal of this study was to examine the link between FC and performance. Because TMS led to changes in both performance and visual network FC, links between behavior and FC should be expected and may shed light on the neural mechanisms of the TMS effect on performance. In addition, it is possible that within-module FC and between-module FC may relate to performance in different ways, as the effect of TMS intensity has been shown in the previous section to interact with FC types (i.e., within- or between-module connectivity). Therefore, the links between visual system FC and task performance were compared in a mixed effect regression model in which the correct response rate was predicted by the between-module and within-module FC of the visual module. The model also accounted for other factors that may affect performance, including TMS intensity, baseline performance, RMT, and gender (see **Fig. S1**). The effects of these factors on performance were assessed using F-contrasts and T-contrasts and are summarized in **Table 1**.

**Table 1.**
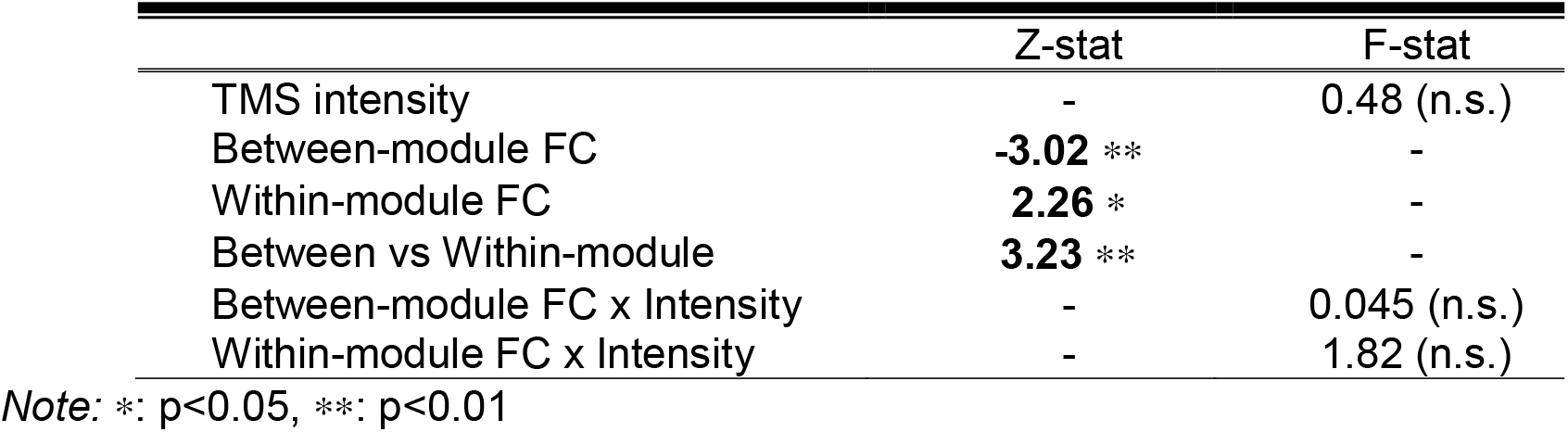
Effects of TMS Intensity (%RMT) and FC type on functional connectivity.

In contrast to the behavioral findings in *Section 3*.*1*, once FC measures were considered in the model, the TMS intensity showed no reliable effect on performance (F_3,64.12_ = 0.48, p = 0.70). In fact, the between-module FC, previously shown to be significantly modulated by TMS, negatively predicted response accuracy (beta = -0.27, z = -3.02, p = 0.0025, **Fig. 5A**). The within-module FC, in contrary, positively predicted response accuracy (beta = 0.12, z = 2.26, p = 0.024, **Fig. 5B**). Such differences in behavioral association were further indicated by a Type by FC interaction (i.e., between-module minus within-module; beta = 0.39, z = 3.23, p = 0.0013), suggesting that the effect of TMS on FC during task performance was dependent on the type of FC. There was no significant Intensity x FC interaction for either within-module FC (F_3, 63.54_ = 1.82, p = 0.15) or between-module FC (F_3, 63.84_ = 0.045, p = 0.99) of the Visual module. In sum, these results point to the network-level functional communication mediating the TMS-behavior relationships.

**Figure 5.**
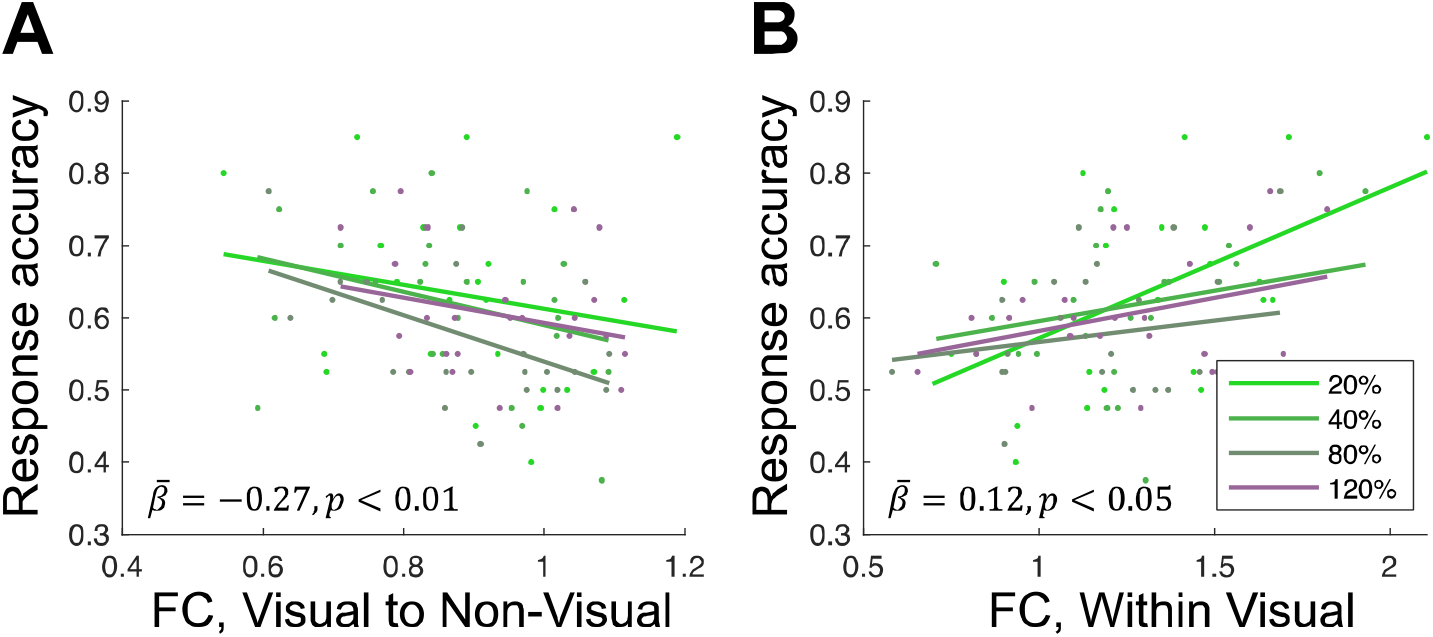
Relationships between task performance and FC either (**A**) between the Visual system and all non-Visual modules, or (**B**) within the Visual network.

#### 3.3.3 Structural constraints in TMS-induced functional connectivity

The third goal of this study was to determine if FC patterns, constrained by the structural connectivity (SC) network, can be altered by TMS. As the pattern of functional communication has been shown to rely on a backbone of white-matter connections (Hermundstad et al., 2013; Goni et al., 2014), it was predicted that the impact of TMS on FC can be reflected by the level of consistency between the functional network and its underlying structural network. At the most basic level, FC-SC similarity can be characterized with Spearman’s correlation between structural and functional networks; implicit in this relatively straightforward approach is the premise that FC should reflect the underlying SC, and that lower FC-SC similarity indicates a greater deviation of the FC pattern from a more original configuration constrained by the structural backbone, hence reflecting a greater disruption of the FC pattern.

For the Visual network, an Intensity by Type repeated measures ANOVA on FC-SC similarity was employed to assess if and how TMS contributes to the change in FC. The analysis revealed a significant effect of Intensity (F_3,63_ = 6.17, p = 9.6e-4), as well as a significant Intensity by Type interaction (F_3,63_ = 4.42, p = 6.9e-3, see **Fig. 6A, 6B**). Separate one-way repeated measure ANOVAs were performed to further examine how between-module and within-module connections were affected. As shown in **Figure 6B**, the TMS intensity effect turned out to be significant for within-module FC-SC similarity (F_3,63_ = 11.18, p = 5.65e-6) in the Visual network, leading to more distinct patterns of FC at higher intensity levels. In comparison, TMS intensity showed no effect for between-module FC-SC similarity (F_3,63_ = 0.65, p = 0.59). Lastly, an exploratory analysis was performed to test whether the FC pattern between other modules could be affected by TMS (**Fig. 6C**), which further confirmed that the TMS-induced disruption of FC pattern was specific to the FC within visual system.

**Figure 6.**
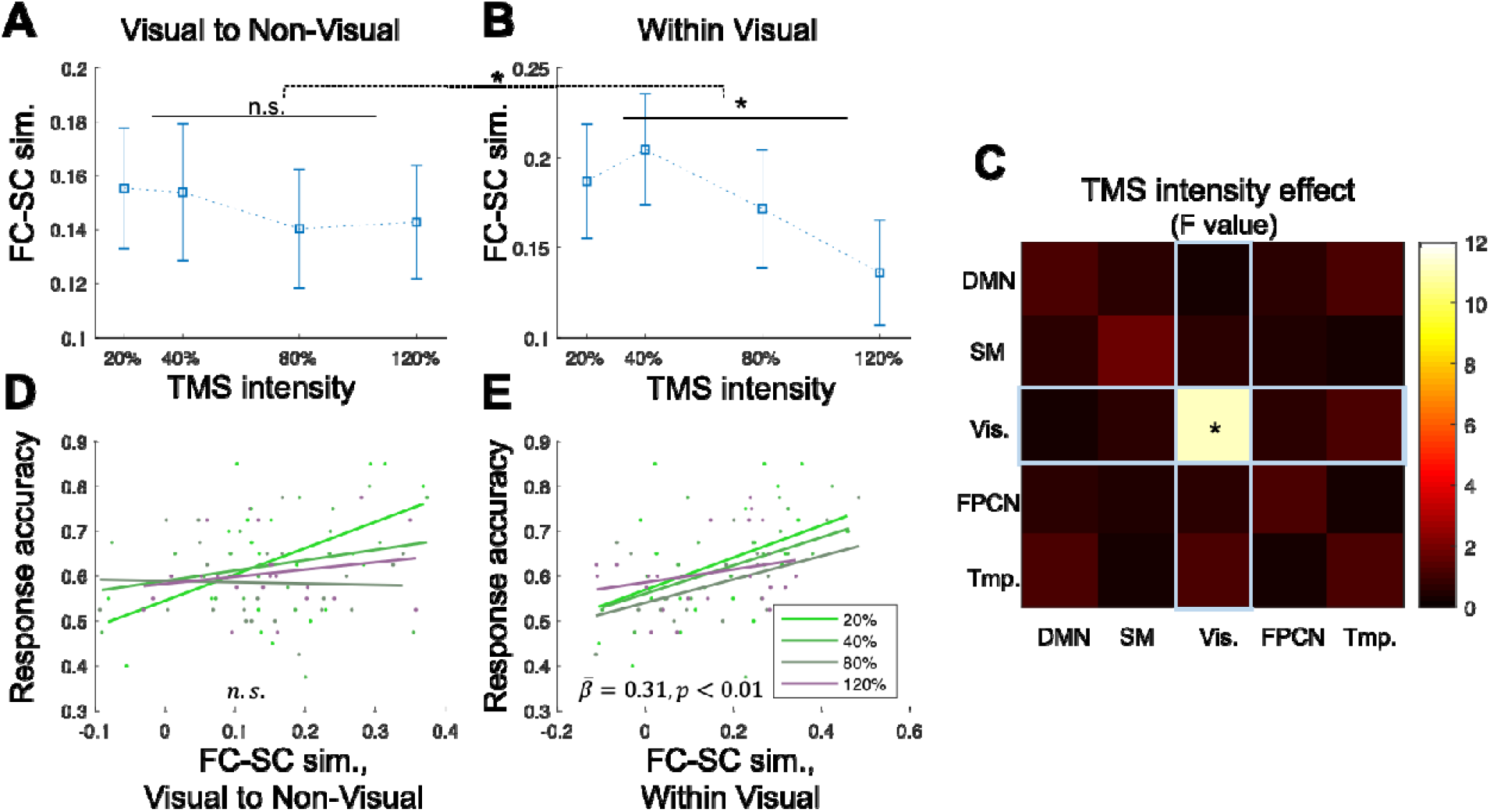
The effects of TMS intensity on visual network FC-SC similarity. Dashed horizontal brace indicates the Intensity x Type interaction; horizontal solid lines indicate main effect of Intensity in the Within Visual **(B)** connections, but not from Visual to Non-Visual **(A)** connections. **C)** The effects of TMS intensity (expressed as F-values) on the patterns of FC (measured as FC-SC similarity) between different modules. **(D)** and **(E)** The relationships between task performance and FC-SC similarity. Error bar indicate standard error of the mean (SEM). Asterisks (*) indicate p<0.05.

Next, to determine if FC-SC coupling influenced motion perception accuracy a mixed effect model was performed in which the correct response rate for each participant, under each TMS intensity level, was assigned as the outcome variable. The TMS intensity levels and its interactions with the between-module FC-SC similarity and the within-module FC-SC similarity of the visual module were modeled as fixed effect. Potential covariates, including sex, coherence of dots, and RMT of each subject, were included in the model and treated as fixed effect as well. The effects of these factors on performance were assessed using F-contrasts and T-contrasts, and they are summarized in **Table 2**. Similar to the previous finding in *Section 3*.*3*.*2*, with the brain measure (FC-SC similarity) taken into account, TMS intensity no longer showed a significant effect on the correct response rate (F_3,56.71_ = 0.19, p = 0.90). The within-module FC-SC similarity of the visual network, previously shown to be modulated by TMS, was positively predicting the correct response rate (beta = 0.31, z = 2.71, p = 0.0068, **Fig. 6E**). The between-module FC-SC similarity of the visual module was not significantly related to correct response rate (beta = -0.04, z = -0.32, p = 0.75, **Fig. 6D**). The Intensity by FC Type interaction effect was found to be trending significant (i.e., between-module minus within-module; beta = 0.36, z = 1.77, p = 0.077).

**Table 2.** Effects of TMS Intensity (%RMT) and FC-SC similarity on Behavior.

|  | Z-stat | F-stat |
| --- | --- | --- |
| TMS intensity | - | 0.19 (n.s.) |
| Between-module FC-SC sim. | -0.32 (n.s.) | - |
| Within-module FC-SC sim. | **2.71** ** | - |
| Between vs Within-module | 1.77 . | - |
| Between-module FC-SC sim. x Intensity | - | 1.66 (n.s.) |
| Within-module FC-SC sim. x Intensity | - | 0.24 (n.s.) |

**Note:** .: $p < 0.1$ ; \*: $p < 0.05$ , \*\*: $p < 0.01$

## 4. DISCUSSION

Current views of the neural effects of TMS are changing. Earlier basic science approaches to TMS in humans focused on the effects of localized stimulation to a specific cortical target. As both imaging and TMS techniques evolved, a more recent focus on bivariate connections from the stimulation site to a subcortical target have revealed promising findings, such as the possibility of targeting subcortical structures with cortical TMS (Wang et al., 2014; Luber et al., in press). However, there is a growing appreciation for how stimulation at a single site can propagate throughout a cortical network (Cocchi et al., 2016) and a growing need to identify which principles govern that flow. Though there are only a limited number of combined TMS-EEG or TMS-fMRI studies considering this problem from a network-level perspective, one general principle that emerges across studies is that local stimulation spreads according to the functional networks associated with the stimulated node (Momi et al., 2021b). This study advances the field by considering this phenomenon during the engagement of an active task, and finds that the effects of TMS on ongoing brain activity are most evident in regions actively engaged with the task at hand. This is a critical extension of expectations of local cortical excitation or inhibition with specific stimulation frequencies (Aydin-Abidin et al., 2008), and to the variability of modulation observed during different brain states (Gollo et al., 2017). By stimulating with bursts of 10 Hz rTMS at different intensities over the primary visual cortex during a motion perception task, and concurrent with fMRI imaging, the present study revealed four novel findings. First, motion processing was selectively impaired at 80% RMT. Second, the connectivity between the visual system and other cortical communities increased with stimulation intensity. Third, intra-community connectivity was beneficial, while the extra-community connectivity induced by TMS was detrimental to performance. Lastly, structural connectivity constrained this effect, such that FC-SC was strongest in low-intensity conditions, and was disrupted with increasing intensity, resulting in poorer performance. These findings are discussed in greater detail below.

Before turning to the connectivity-related findings, it is important to consider the observation that TMS-related impairment of motion discrimination was selective to the 80% RMT condition. No differences in discrimination accuracy were seen between the highest (120% RMT) and lowest (20% RMT) stimulation intensities, suggesting that the disruptive effects were attributable to a neural response in visual cortex, and not distraction-related confounds associated with increasing TMS intensity (e.g., noise, cutaneous sensation). Past studies have reported a wide variety of behavioral effects that result from single or paired pulse TMS over a range of factors, such as timing, intensity, and the ongoing activation state of the stimulated region. For example, in primary motor cortex the threshold for inhibitory effects of TMS is approximately 80% of RMT (Kallioniemi et al., 2014). Thus, TMS intensity can determine the local balance between inhibitory and excitatory circuit recruitment. In another example, single pulses of 80% RMT TMS timed to the onset of the N2 latency of MT+ facilitate motion perception (Gamboa Arana et al., 2020). In addition to the possibility of a specific balance of inhibitory and excitatory effects, such findings may be explained by a stochastic resonance account, such that the strength of a stimulus can be improved by externally enhancing the ongoing neural activity. Despite a similar task paradigm and dosing structure, the current study instead revealed a disruption of performance at 80% RMT, perhaps due to the differences in cortical target (V1 vs. MT+), or the use of three pulses (spaced at 10Hz) instead of one pulse of TMS. Furthermore, this single pulse was timed to the onset of a specific, motion sensitive ERP. Nonetheless, the current results may abide by the principle that a selective modulation of performance is based on the engagement with ongoing brain activity, such that that 10 Hz pulses timed to the onset of motion disrupted the downstream flow of information from primary visual cortex to motion-sensitive cortex (MT+). This pattern of results also lends support to a more global model of the effects of TMS on behavior (Silvanto and Cattaneo, 2021) that posits distinct facilitatory and suppressive ranges as a function of TMS intensity, and which are shifted by state-dependent changes in neural excitability (e.g., during stimulus processing vs. rest states).

Turning to the general connectivity changes in response to TMS, the current study first found that local connectivity within the visual system was more resilient to stimulation intensity than global connectivity from the visual system to the rest of the brain. Such a result is consistent with the notion that the network responsible for visual motion is an optimized system that operates best when allowed to segregate from other cortical networks. In contrast, when stimulation intensity induced an increased spread of connectivity between modules, this more global pattern had negative consequences for behavior. In fact, when we examined the source of these intensity-related increases in between-module FC, we found greater FC with the somatosensory-motor and temporal networks, consistent with the idea that increasing intensity of TMS elicited greater network activity due to increasing motor (SM) or auditory (Temporal) sensation. Thus, an important consideration for studies examining the network-level effects of TMS is to consider how some of these distant effects can be attributed to the sensory properties of TMS stimulation, as revealed by conditions that mimic the sound and somatosensory attributes of TMS while minimizing intracranial E-fields (“sham” conditions). In some cases, these confounds may be unavoidable (see Conde et al., 2019 for innovative post-processing techniques) but they are not necessarily a barrier to interpreting effects in non-sensorial regions.

An interesting, if overlooked, observation in many concurrent TMS-fMRI studies is that the effects of localized TMS are not strongest at the site of stimulation and are often seen in distant regions of the cortex. While some of these distant effects can be attributed to the sensory properties of TMS stimulation, discussed above, a recent meta-analysis revealed that this surprising absence of a local effect generalizes across many different stimulation paradigms (Rafiei and Rahnev, 2021). More broadly, it is important to consider how such networks may be influenced or modulated by the level of attention associated with increasing TMS intensity. Selective attention can modulate the ongoing processing in both visual (Ballesteros et al., 2008) and motor (Johansen-Berg and Matthews, 2002) regions, and thus it is possible that heightened responses in sensory regions reflect an increase in the alerting or arousal effect associated with increasing TMS intensity, in addition to the input from more basic sensory afferents. While not an explicit focus of this study, such an interpretation is not inconsistent with the finding that attention-driven MT+ activity during motion-perception tasks is influenced by the gating of V1 and V5 connectivity by the posterior parietal cortex (Silvanto et al., 2005). Taken together, this perspective links how a network-level interpretation of TMS effects can better explain the function of single region in a global perspective. While there are many studies on the impact of TMS on local function of the visual regions during motion paradigms (Campana et al., 2002; Cowey et al., 2006; Amemiya et al., 2017), there is minimal research on the network-based influence on this region.

Another important finding was that connectivity *within* the visual network was positively associated with motion accuracy, whereas the connectivity between visual and other networks was negatively associated motion accuracy. Furthermore, follow-up tests demonstrated that the positive correlation between within-module FC and response accuracy was strongest at the lowest TMS intensity, when our exogenous stimulation was unlikely to have much of an effect on cortical excitability. In contrast, increasingly worse performance was associated with more global connectivity at higher levels of TMS intensity (**Table 1**). Such a benefit to more segregated cortical networks is well known in developmental (Stevens et al., 2012) and aging (Betzel et al., 2014; Crowell et al., 2020) literatures, but this is the first finding to suggest an active, parametric perturbation of cortical modularity and its consequences for behavior. An alternative explanation for these effects is that greater stimulation intensity also activated auditory and somatosensory-motor regions, and that this influence of non-visual systems resulted in more interference to the efficient processing of visual stimuli. However, this view is not supported by the fact that motion discrimination performance was greater at 120% RMT than 80% RMT, suggesting a non-linear relationship between delivered E-field and its effect on behavior in this system.

Lastly, an analysis of SC data revealed that the propagation of these effects was largely constrained by multiple indices of white matter anatomy. Two related findings support the notion that the FC patterns associated with localized TMS stimulation are constrained by structural architecture. First, we examined how FC-SC correlations change as a function of TMS intensity. Collectively, significant effects of both Intensity and a significant Intensity × FC Type interaction, as well as the decline in within-module connectivity with increasing intensity all suggest that TMS disrupted “normal” visual network functioning and introducing exogenous stimulation disrupted successful motion processing by decoupling FC and SC patterns. Second, and supporting this inference, an Intensity × FC Type Repeated Measures ANOVA revealed a pattern that mirrors the FC findings above, such that the coupling between structural and FC predicted task performance for the local within-module regions, but more global between-module couplings had no influence on motion perception (**Table 2**). In our study the FC-SC coupling ranged between r = [0.0–0.3], consistent with other many other investigations between functional and structural network data relying on one-to-one correspondence between FC and SC connections (Honey et al., 2007; van den Heuvel et al., 2009; Horn et al., 2014). This suggests that the upper bound of most FC-SC correlations is not very high (Suarez et al., 2020). Future approaches to characterizing FS-SC relationships may consider the value of derived properties of individual nodes that describe their global influence, including modal controllability—which purports to describe the energy associated with transitioning between brain states (Gu et al., 2017; Beynel et al., 2020b).

While the current study represents a novel application of concurrent TMS-fMRI to answer unexplained phenomena describing the spread of focal neurostimulation across the cortex, several limitations were encountered in the pursuit of this goal. First, the study design did not incorporate sham conditions that closely replicate the sound and scalp sensation of TMS without the delivery of significant E-field to the brain. Nevertheless, the results of the parametric investigation of TMS intensity at the target support a cortical origin of the stimulation effects. Second, because the goal of the current study was to investigate multiple stimulation intensities, it was not feasible to complete within a typical MRI scanning session to assess a complementary stimulation site to examine the potential dissociations between task-relevant and task-irrelevant cortical sites. Lastly, the current design relied on a trial-related timing of 10 Hz pulses concurrent with the task; these design choices represent only one of a number of parameters for frequency, number of pulses, and timing relative to the motion perception task. While the prolonged effects of rTMS are generally assumed to relate to synaptic plasticity, the detailed effects of stimulation on a region’s excitability via long-term depression or potentiation remains a matter of investigation.

The observed pattern of findings suggests that inducing a more global pattern of connectivity with higher intensity TMS contributes to disruptions in visual motion perception. Our results show that this spread is largely limited to the network of cortical sites involved in the active task (i.e., visual motion), and that TMS-related disruptions of this network are detrimental to cognitive performance. Lastly, we find that the functional connectivity spread with increasing TMS is largely constrained by structural connectivity, suggesting that the use of structural network information can help to predict at least some of the influence of local perturbations on a more global cortical network. Thus, while it is widely known that TMS may disrupt cognition, these results help us to explain *how* TMS disrupts local and global networks to induce perceptual change.

## ACKNOWLEDGMENTS

We thank Lamont Conyers and Jennifer Graves for extensive MRI support, and all subjects for their participation. This study was supported by the National Institute of Health, Brain Initiative 1RF1- MH114253, R21-AG058161, and K01-AG053539.

